# Circadian rhythms in bipolar disorder patient-derived neurons predict lithium response

**DOI:** 10.1101/2020.12.14.422616

**Authors:** Himanshu K. Mishra, Noelle M. Ying, Angelica Luis, Heather Wei, Metta Nguyen, Timothy Nakhla, Sara Vandenburgh, Martin Alda, Wade H. Berrettini, Kristen J. Brennand, Joseph R. Calabrese, William H. Coryell, Mark A. Frye, Fred H. Gage, Elliot S. Gershon, Melvin G. McInnis, Caroline M. Nievergelt, John I. Nurnberger, Paul D. Shilling, Ketil J. Oedegaard, Peter P. Zandi, The Pharmacogenomics of Bipolar Disorder Study, John R. Kelsoe, David K Welsh, Michael J. McCarthy

## Abstract

Bipolar disorder (BD) is a neuropsychiatric disorder with genetic risk factors defined by recurrent episodes of mania/hypomania, depression and circadian rhythm abnormalities. While lithium is an effective drug for BD, 30-40% of patients fail to respond adequately to treatment. Previous work has demonstrated that lithium affects the expression of “clock genes” and that lithium responders (Li-R) can be distinguished from non-responders (Li-NR) by differences in circadian rhythms. However, rhythm abnormalities in BD have not been evaluated in neurons and it is unknown if neuronal rhythms differ between Li-R and Li-NR. We used induced pluripotent stem cells (iPSCs) to culture neuronal precursor cells (NPC) and glutamatergic neurons from BD patients and controls. We identified strong circadian rhythms in *Per2-luc* expression in NPCs and neurons from controls and Li-R. NPC rhythms in Li-R had a shorter circadian period. Li-NR rhythms were low-amplitude and profoundly weakened. In NPCs and neurons, expression of *PER2* was higher in both BD groups compared to controls. In neurons, PER2 protein expression was higher in BD than controls, especially in Li-NR samples. In single cells, NPC and neuron rhythms in both BD groups were desynchronized compared to controls. Lithium lengthened period in Li-R and control neurons but failed to alter rhythms in Li-NR. In contrast, temperature entrainment increased amplitude across all groups, and partly restored rhythms in Li-NR neurons. We conclude that neuronal circadian rhythm abnormalities are present in BD and most pronounced in Li-NR. Rhythm deficits in BD may be partly reversible through stimulation of entrainment pathways.

## Introduction

Bipolar disorder (BD) is a psychiatric illness characterized by recurrent episodes of mania/hypomania, depression and increased risk for suicide^1^. With heritability estimates of 70– 85%, it is clear there is a genetic component to BD, but the biological origins remain incompletely characterized ^2, 3^. Lithium responsiveness has been proposed as a genetically-influenced clinical subphenotype that may help organize the heterogeneity of BD^4, 5^. Besides mood disturbances, BD is associated with altered rhythms in sleep and activity. This observation underlies the hypothesis that circadian rhythm disruption is a component of the illness ^6^ and that assessing circadian rhythm disruption in BD may be another strategy for reducing heterogeneity ^7^. Previously, we found overlap between lithium-response and circadian rhythms in BD, showing that BD patients with high levels of morningness were more lithium responsive. Moreover, fibroblasts cultured from lithium-responsive BD patients had shorter circadian periods compared to cells from lithium non-responders ^8^. Circadian rhythms are cell-autonomous ∼24 h oscillations generated from a network of ∼20 ‘clock genes’ that coordinate timing through transcriptional/translational feedback loops ^9^. The CLOCK/BMAL1 complex drives the expression of PER1/2/3 and CRY1/2 to provide inhibitory feedback. In mice, the CLOCKΔ19 mutant has a mania-like phenotype ^10^, while BMAL1 knockdown in the suprachiasmatic nucleus (SCN) models depression ^11^. However, animal models fail to capture key features of BD, and human cellular models remain essential to understanding biological mechanisms. Previous studies have shown that molecular rhythm abnormalities can be measured in BD patient fibroblasts using bioluminescent reporters ^8, 12, 13^. However, the fibroblast model lacks essential features, and neurons may have advantages in modelling cellular mechanisms underlying neuropsychiatric disorders like BD.

Technological advances enable the use of induced pluripotent stem cells (iPSCs) to grow human neurons *in vitro*, providing a powerful tool for investigating the neurobiology of psychiatric disorders. Previous work has shown that iPSC-derived BD neurons showed widespread gene expression alterations and electrical hyperexcitability that was selectively reduced by lithium in neurons from lithium responders (Li-R) ^14^. Presently, we used a similar approach to analyze circadian rhythms in iPSC-derived neuronal progenitor cells (NPCs) and neurons derived from Li-R and Li-NR BD patients. We observed profound circadian rhythm disruptions in BD cells, especially in Li-NR. These observations further implicate the circadian clock in BD, and may provide the basis for novel approaches to study the mechanisms underlying the disorder.

## Methods

### Human subjects

Subjects provided written informed consent in accordance with all relevant regulations. BD subjects (n=5) were males of European ancestry, aged 22-69 y, diagnosed as BD type I (Table S1). Subjects were assessed for clinical response to lithium monotherapy in a prospective clinical trial (supplemental methods) as reported previously ^5^. BD donors were selected based upon their clinical response to lithium, identifying subjects from the tails of the distribution to enrich for the most and least lithium responsive. Age, sex and ancestry-matched controls (n=4) were determined to be free of a psychiatric or substance use diagnosis by clinical interview. Skin biopsies from the deltoid region were used to culture fibroblasts using standard procedures. Cultures were then frozen until use for iPSCs ^14^.

### Human iPSC cultures

IPSCs were generated as described ^14^. Cyto-Tune Sendai reprogramming kit (ThermoFisher) was used according to the manufacturer’s instructions and colonies were expanded on Matrigel-coated (BD Biosciences) plates in mTeSR1 medium (Stemcell Technologies). Lack of genomic integration of Sendai virus into iPSCs was confirmed by PCR and karyotype analysis confirmed the expected chromosomal copy numbers ^15^. Every 7– 8 days, iPSCs were treated with collagenase, and passaged. To verify pluripotency, cells were characterized for Nanog and Tra1-60 ^16^.

### Generation of neural progenitors and neurons

Colonies were transferred to ultra-low attachment plates (Corning) to generate embryoid bodies, EBs ^16^. EBs were treated with neural induction medium [NIM: DMEM/F12/glutamax/N2/B27 (Invitrogen) and pen/strep)] for 1 week. Cells were transferred to polyornithine/laminin (Invitrogen)-coated plates. After 7 d, neural rosettes were dissected from the EBs and expanded in NIM containing 20 ng/ml FGF2 (Preprotech). Rosettes were dispersed to form NPCs and passaged every 5-7 d. To obtain mature neurons, NPCs were treated with NIM with 20 ng/ml BDNF (Preprotech), 20 ng/ml GDNF (Peprotech), 1 mm dibutyryl-cyclic AMP (Sigma–Aldrich) and 200 nm ascorbic acid (Sigma– Aldrich), and cultured with weekly half media changes. Neurons used in experiments were differentiated for 6-8 wk. At least two clones were independently grown for each line. In some instances, it was not possible to grow sufficient numbers of cells for all experiments. A detailed accounting of cells allocated for each experiment is provided (Table S2-S3).

### Measurement of circadian rhythms

Rhythms in NPCs (n=4 control, 2 Li-R, 3 Li-NR) and neurons (n=3 control, 2 Li-R, 3 Li-NR) were measured using the *Per2-luc* bioluminescent reporter ^12^. Cells were cultured in triplicate 250 x10^3^/35 mm plate then infected with lentiviral *Per2-luc* for 48 h. Prior to recording, cells were synchronized with 10 µM forskolin (Tocris) for 2 h. Following a media change, photoemissions were recorded for 70 s every 10 min for 5-7 d using a luminometer (Actimetrics). Detrended data were fit to a damped sine wave and rhythm parameters were calculated using LumiCycle Analysis (Actimetrics). In some luminometer studies, rhythms were measured in cultures treated with lithium (1-10mM) or vehicle.

### Gene expression assays

For gene expression analyses, NPCs (n=2 control, 2 Li-R, 2 Li-NR) or neurons (n=2 control, 2 Li-R, 3 Li-NR) were grown to a density of 400×10^3^/well. Cells were synchronized using 10 µM forskolin (Tocris). After 16 h, plates were collected at 6 h intervals over 24 h. Plates were washed with cold PBS and frozen at −80°C. RNA was extracted using RNeasy kit (Qiagen) and quantified by spectrophotometer. RNA (500 ng) was reverse transcribed into cDNA using QuantiTect RT-PCR kit (Qiagen). RT-PCR was performed on a BioRad CFX384 thermocycler using pre-validated Taqman primers (Applied Biosystems) for *BMAL1 (ARNTL), CLOCK, CRY1, NR1D1, PER1/2/3*, and *RORA*. Expression was normalized to *GAPDH*, a non-rhythmic reference gene ^17^ and normalized to mean expression over 24 h. Using data matched for sample and experiment, correlation coefficient across time for each gene pair were calculated to identify network-level patterns in expression. Correlation coefficients were z-transformed and network connectivity was visualized using Cytoscape ^18^ and analyzed by 2-way ANOVA.

### Single-cell imaging in NPCs and neurons

Imaging was performed using *Per2-luc* ^19, 20^. *Per2-luc* transfected NPC/neurons (90×10^3^ cells/well) were synchronized with forskolin 10 µM (Tocris) for 2 h. Cells were placed upon an inverted microscope (Olympus) and maintained in darkness at 36°C. Light was collected by an Olympus XLFLUOR 4x objective and camera (Series 800, Spectral Instruments). Images were collected every 30 min for 4-5 days and analyzed with MetaMorph (Universal Imaging). Rhythmic cells were defined as having a period in the circadian range of 20-28 h, with the percent of fast Fourier transform (FFT) spectral power in a 1 Hz window ≥15% for NPCs and ≥ 8% for neurons.

### Immunocytochemistry

Cells on polyornithine/laminin coated coverslips were fixed with 4% paraformaldehyde, then permeabilized using 0.2% TritonX-100 (Sigma). Non-specific binding was blocked using 5% donkey serum (Jackson ImmunoResearch). Primary antibodies (Table S4) diluted in 2% donkey serum were added to coverslips and incubated overnight at 4°C. Coverslips were washed with 0.2% TritonX-100, and incubated with secondary antibodies (1:500) diluted in 2% donkey serum for 2 h at 20°C. Coverslips were incubated in DAPI (ThermoFisher) for 5 min, mounted, and visualized using a fluorescent microscope (Leica).

### Statistical analysis

Given the scarcity of samples, statistical power was limited. To increase power, each clone and corresponding technical replicate/single cell was considered independently which accurately considers variance related to passage number, culture conditions, *Per2-luc* infection and unattributable experimental error, but may underestimate variance between subjects. Efforts were made to balance replicate numbers to avoid over-sampling any particular donor. ANOVAs, t-tests and chi-squares were performed using GraphPad Prism 5.0. Circular statistics were analyzed using Oriana 4.0. Significance was defined by p-value ≤ 0.05. Data are displayed as mean ± standard error of mean (SEM).

## Results

### Characterization of neuronal cultures

As expected, iPSCs expressed pluripotency markers Nanog and Tra-1-60 (Figure 1B). NPCs expressed NESTIN and SOX2 (Figure 1C). Approximately 70-80% of differentiated cells expressed β-III tubulin (TUJ1, Figure 1D) while 10-20% expressed GFAP (Figure S1). Approximately ∼80% of TUJ1-stained neurons were positive for vesicular-glutamate transporter 2 (VGLUT2), indicating enrichment of excitatory, glutamatergic neurons (Figure 1E and S1).

**Figure 1:**
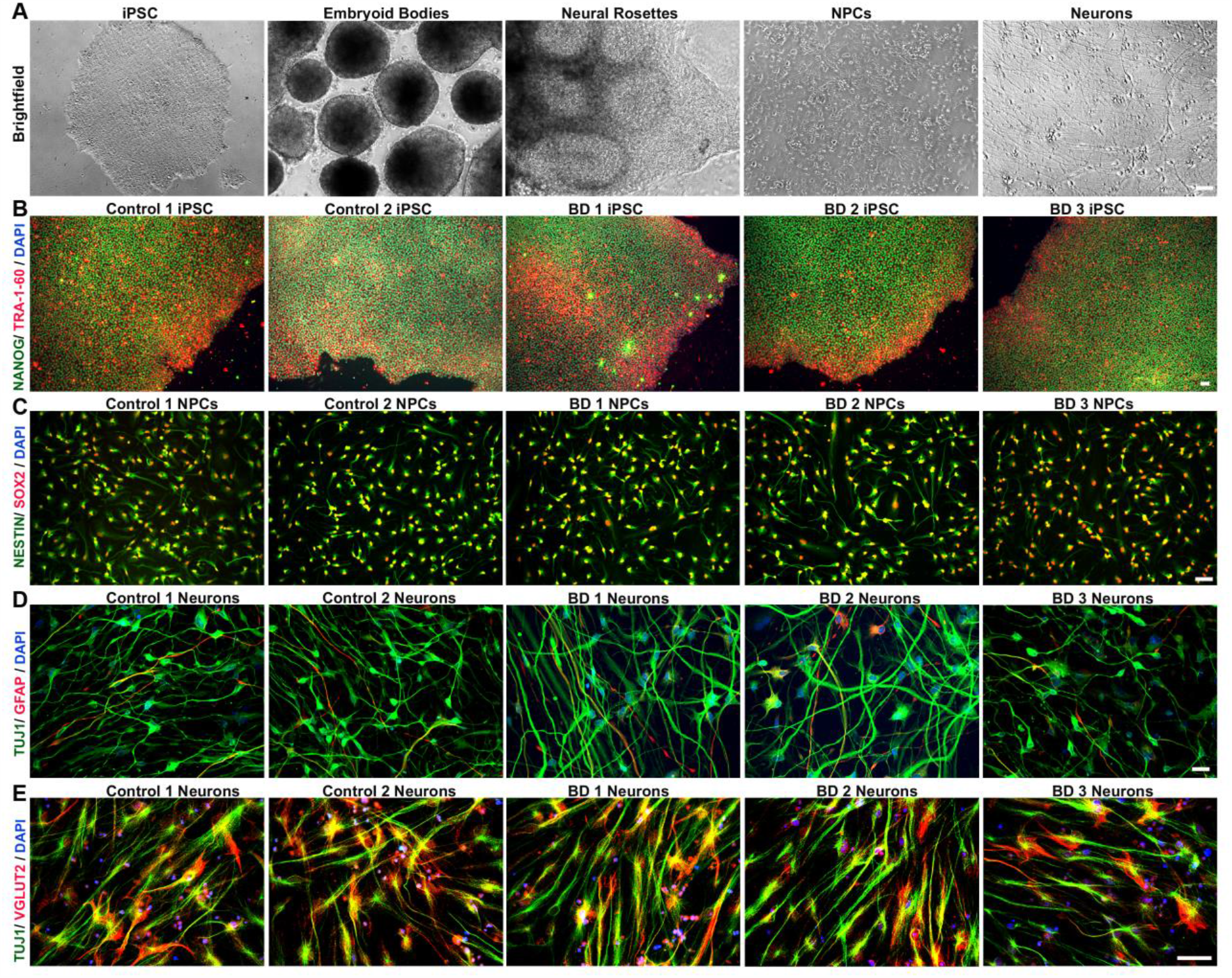
Generation of human neurons from induced pluripotent stem cells. Control and BD iPSCs are morphologically similar, characterized by well-defined borders, circular shape and expression of pluripotency markers. (A) Brightfield images showing stages of embryoid bodies based neural differentiation paradigm for iPSCs. (B) Representative image of an iPSC clone co-labeled with pluripotency markers, TRA 1-60 (red), and NANOG (green). (C) NPCs generated from control and BD iPSCs co-express early cortical neural precursor markers NESTIN (green) and SOX-2 (red). (D) Differentiated neurons expressing the neuron-specific marker TUJ1, and only weakly express the glia-specific marker GFAP. (E) Representative images of control and BD neurons showing widespread co-localization of vesicular-glutamate transporter 2 (VGLUT2) in TUJ1 positive cells, indicating enrichment of glutamatergic neurons. Cell nuclei are stained with DAPI (B-E). Antibodies used are listed in Table S2. Scale bars = 50µm. BD: bipolar disorder; DAPI: 4’,6-diamidino-2-phenylindole; GFAP: glial fibrillary acidic protein; iPSCs: induced pluripotent stem cells; NPCs: neural progenitor cell; TUJ1: β-III tubulin; VGLUT2: vesicular-glutamate transporter Numbers in the titles were arbitrarily assigned for identification.

### Circadian rhythms in NPCs and neurons

To model neurodevelopmental aspects of BD, we used both NPCs and neurons in circadian rhythm experiments. Consistent with our previous studies in fibroblasts ^8, 12^, we observed robust *Per2-luc* rhythms in NPCs and neurons (Figure 2A, G). Control NPCs showed high amplitude rhythms that persisted for >5 days. In BD NPCs, Li-NR samples exhibited weak, low-amplitude rhythms, while Li-R rhythms were similar in amplitude to controls (mean amplitude counts/sec ± SEM control = 0.06 ± 0.01, Li-NR = 0.03 ± 0.003, Li-R = 0.06 ± 0.01, p<0.005, n=4 control, 2 Li-R, 3 Li-NR, Figure 2 A-F). Period was significantly shorter in Li-R compared to controls but not significantly different in Li-NR (mean period h ± SEM: control = 25.98 ± 0.32, Li-R = 21.85 ± 0.44, Li-NR = 24.70 ± 0.50, p < 0.001, n=4 control, 2 Li-R, 3 Li-NR, Figure 2E). Rhythm damping was faster in NPCs from Li-NR (mean damping constant, days ± SEM: control = 3.40 ± 0.66, Li-R = 2.59 ± 0.76, Li-NR = 1.54 ± 0.17; p<0.005, n=4 control, 2 Li-R, 3 Li-NR, Figure 2F). In neurons, there were no significant differences in period among the groups (mean period h ± SEM: control = 25.65 ± 0.22 h, Li-R = 25.60 ± 0.27, Li-NR = 25.49 ± 0.37; n=3 control, 2 Li-R, 3 Li-NR, Figure 2K). Damping was also faster in neurons from Li-NR (mean damping constant, days ± SEM: control = 1.832 ± 0.11, Li-R = 2.41 ± 0.34, Li-NR = 1.34 ± 0.31; n=3 control, 2 Li-R, 3 Li-NR, Fig 2L).

**Figure 2:**
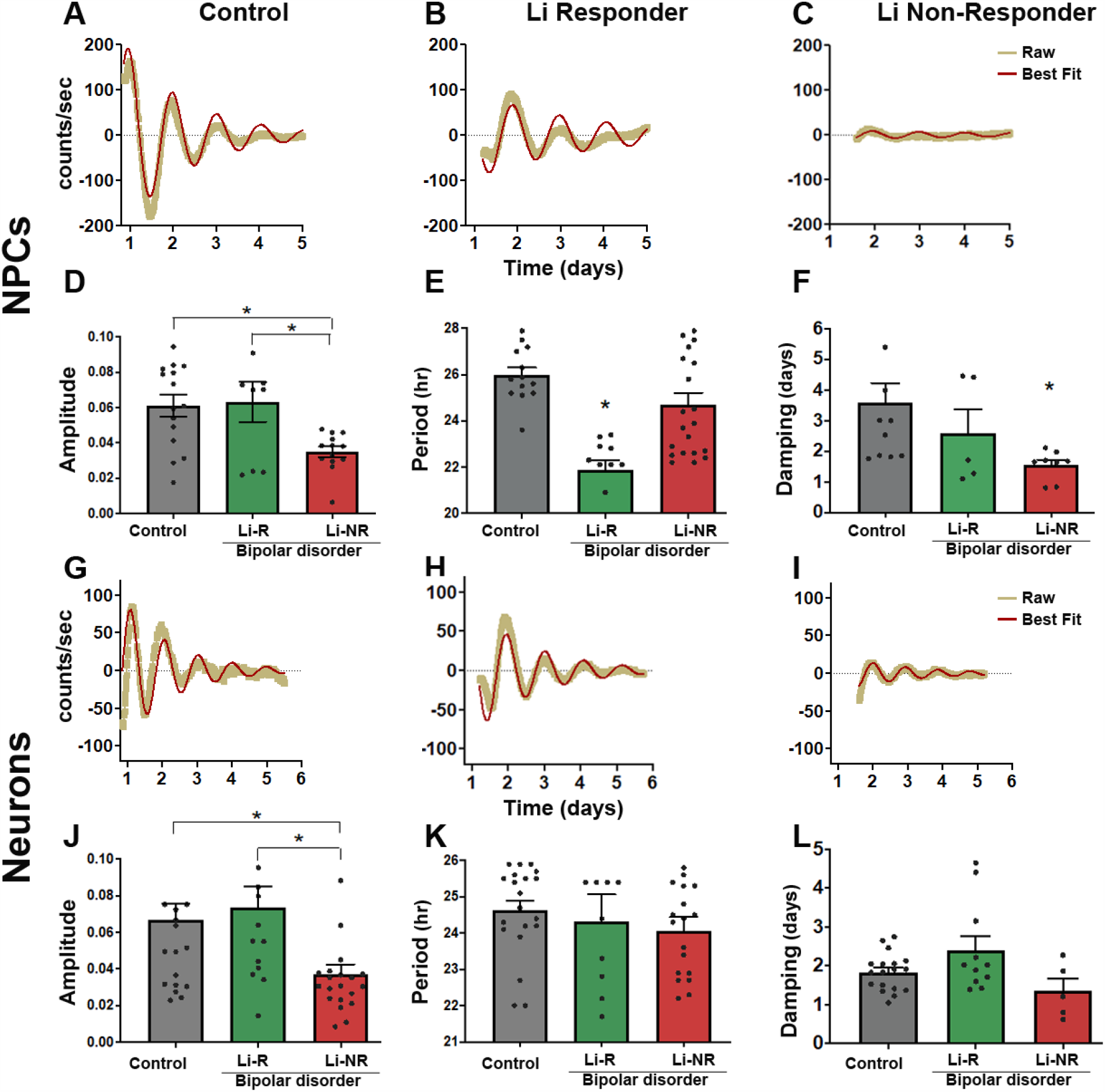
Luminometer assays of circadian rhythms in NPC and neurons. **(A-C)** Representative traces of Per2-luc rhythms measured by luminometer in plates of NPCs from **(A)** controls **(B**) lithium responders (Li-R), and **(C)** lithium non-responders (Li-NR). Yellow indicates raw counts, red indicates best fit line. **(D)** Normalized rhythm amplitudes (with units corrected for brightness) were similar in control and Li-R cells, but significantly lower in NPCs from Li-NR (one-way ANOVA p<0.005, * indicates Li-NR lower in post-hoc test). **(E)** Li-R NPCs had shorter rhythm periods compared to controls and Li-NR (one-way ANOVA, p < 0.004). **(F)** Rhythm damping was faster in NPCs from Li-NR (one-way ANOVA p<0.005, * indicates Li-NR lower in post-hoc test). **(G-I)** Representative traces of *Per2-luc* rhythms in neurons from **(G)** control, **(H)** Li-R and **(I)** Li-NR. Yellow indicates raw counts, red indicates best fit line. **(J)** Normalized amplitude was significantly lower in neurons from Li-NR vs. control and Li-R (one-way ANOVA, p < 0.005). **(K)** No significant difference was observed in rhythm period in neurons from control, Li-R and Li-NR donors. **(L)** Rhythm damping (expressed as damping constant) was faster in neurons from Li-NR (one-way ANOVA, p < 0.05). NPC data reflect the results of n=4 control, 2 Li-R, and 3 Li-NR cell lines, recorded in triplicate, repeated in three separate experimental trials. Neuron data reflect the means of n=3 controls, 2 Li-R, and 3 Li-NR cell lines, recorded in triplicate, repeated across five separate experimental trials. Statistically significant differences p<0.05 are indicated by *. Error bars indicates standard error of the mean (SEM).

### Circadian rhythms in single-cells

We next conducted single-cell bioluminescent imaging studies. In NPCs, there was no significant difference in the proportion of rhythmic cells between controls vs. BD (% rhythmic NPCs: control=22, Li-R=20, Li-NR=11, p=0.13). Interestingly, in controls, differentiated neurons became significantly more rhythmic than NPCs, suggesting that healthy neurons become more rhythmic upon maturation (control neuron=37% vs. NPC=22%, p=0.01). In contrast, neither group of BD neurons was more rhythmic than the corresponding NPC sample (Li-R neuron=25% vs. NPC=20%; Li-NR neuron 12% vs. NPC=11%, both p>0.05). Consequently, in neurons the proportion of rhythmic cells was significantly higher in controls than in BD (control 37% vs. Li-R 25% vs. Li-NR 12%, Figure 3B). To determine the strength of rhythms in individual cells, we measured spectral power in the circadian range. In NPCs, Li-R cells had significantly weaker rhythms compared to Li-NR and controls (mean FFT ± SEM: control: 0.24 ± 0.01 vs. Li-R: 0.18 ± 0.01, Li-NR 0.25± 0.01, Figure 3C). In neurons, we again found evidence of weaker rhythms in BD (mean FFT ± SEM: control= 0.08 ± 0.01 vs. Li-R 0.05 ± 0.01, vs Li-NR 0.04 ± 0.01 Figure 3D). Restricting the analysis to rhythmic cells, we analyzed phase. In both NPCs and neurons, controls showed significant phase clustering, whereas in the two BD groups (both Li-R and Li-NR), phase was randomly dispersed across the circadian cycle (NPCs: Figure 3E-I, Neurons: Figure 3F-L).

**Figure 3:**
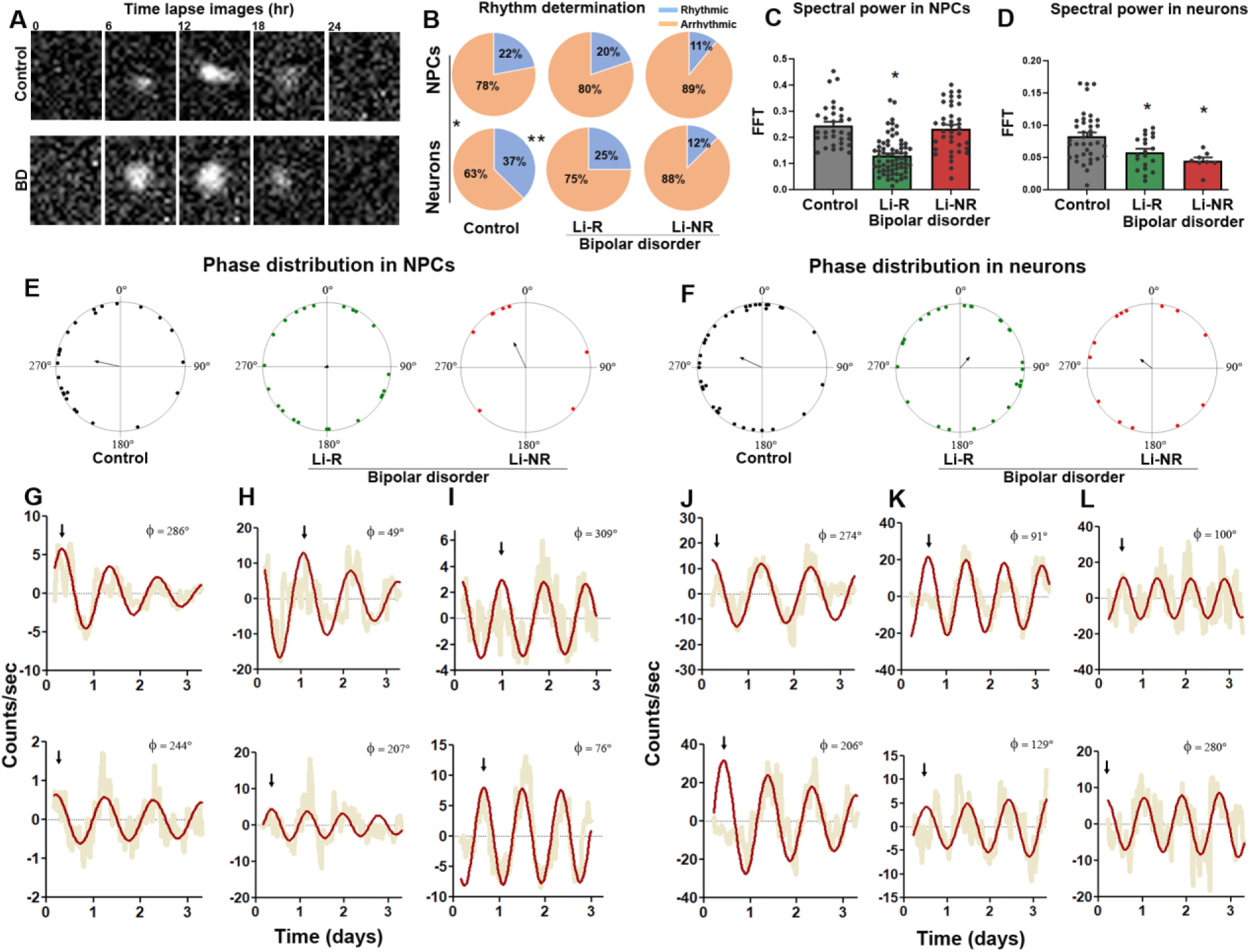
Single-cell assays of circadian rhythms in NPC and neurons. **(A)** Representative time-series images of *Per2-luc* rhythms in single neurons from a control and BD donor. **(B)** Pie charts describing the proportion of cells meeting the definition of rhythmic from control, lithium responder (Li-R) and lithium non-responder (Li-NR) donors. In control samples, the proportion of rhythmic cells increased in differentiated neurons vs. neural progenitor cells (NPCs) (control: χ2 = 6.1(1), p<0.05, indicated by * and vertical line). In cells from Li-R and Li-NR, NPCs and neurons were rhythmic in similar proportions suggesting a lack of developmental effect on the circadian clock in these BD samples [Li-R: χ2=0.54 (1), p=0.46, Li-NR: χ2=0.04(1), p=0.84]. In NPCs, the proportion of rhythmic cells was similar across controls, Li-R and Li-NR [χ2=4.02 (2), p=0.13]. In neurons, there were significantly more rhythmic cells in control vs. Li-R and Li-NR samples [χ2 = 6.23 (2), p < 0.05 indicated by **]. **(C, D)** Single-cell rhythm strength, as measured by relative spectral power in the circadian range. In Li-R, both NPCs and neuron rhythms were weaker vs. controls. In Li-NR, while the total number of rhythmic NPCs was low, those that were detected had normal rhythm strength, whereas rhythmic neurons from Li-NR had significantly weaker rhythms (Analyses by one-way ANOVA revealed for NPCs: F=6.62 p<0.005; neurons: F=6.58 p<0.005. For NPC: N=60-80 cells/line from n=2 control, 2 Li-R, and 1 Li-NR donor, for neurons: N=60-80 cells/line from n=3 control, 2 Li-R, and 1 Li-NR donor). Single-cell analyses indicate significant circadian phase clustering in control NPCs **(E)** and neurons **(F)** (Rayleigh test, p = 0.01), but not in NPCs or neurons from Li-R or Li-NR subjects (p>0.05). Phase is shown using polar coordinates, with dots indicating circadian phases of individual cells. Because phase is undefined in non-rhythmic cells, only rhythmic cells are plotted, and the number of plotted cells is therefore smaller for Li-NR NPCs and neurons. Arrows indicate the mean phase vector for each group. Example pairs of rhythm traces from control, Li-R, and Li-NR NPCs **(G-I)** or neurons **(J-L)** are shown, illustrating representative phase relationships among single-cells from each group (compare top vs. bottom). Yellow indicates raw counts, red indicates best fit curve. Φ indicates phase value (0-360°). Arrows indicate times of first peaks, to facilitate visual comparisons of phase between traces.

### Clock gene expression in NPCs and neurons

We next conducted studies of gene expression in NPCs and neurons. In both cell types, we found that expression of *PER2* was significantly higher in both BD groups vs. controls, (Fig 4A-J, for NPC: n=3 control, 2 Li-R, 2 Li-NR, for neurons n=3 control, 2 Li-R, 3 Li-NR). *PER2* expression was increased by 2-4 fold in both Li-R and Li-NR (Figure 4A, 4F). In NPCs, *CRY1* expression differed significantly over time, but showed no significant group difference. In neurons, *CRY1* was significantly increased by in Li-R neurons compared to controls and Li-NR (Fig 4B, 4G). In NPCs, *BMAL1, CLOCK, RORA, CRY2* and *NR1D1* expression was similar in BD and controls (Figure 4C-E, S4A-B). In neurons, *BMAL1* expression was significantly increased in the Li-R cells compared to controls (Figure 4I, 4J), while there were no differences in *CLOCK, CRY2, NR1D1*, or *RORA* (Figure 4G, S4C-D). We examined the clock gene expression networks in neurons, and found the control and BD (combined Li-R and Li-NR) networks differed significantly, with the greatest differences in the coordination between *BMAL1-CLOCK* and *BMAL1-PER2* (Figure 4K). Using immunolabelling, we found widespread PER2 staining in the neurites of TUJ1-positive neurons. PER2 expression in neurons was significantly higher in both BD groups vs. controls (mean immunoreactivity ± SEM: control: 15.25 ± 0.34 vs. Li-R 17.24 ± 0.72 vs. Li-NR 27.46 ± 0.88, p<0.0001, n=3 control, 2 Li-R, 3 Li-NR donors, Figure 4L-M, S4E-F). Post-hoc analyses revealed that PER2 protein was also significantly increased in Li-NR neurons compared to Li-R.

**Figure 4:**
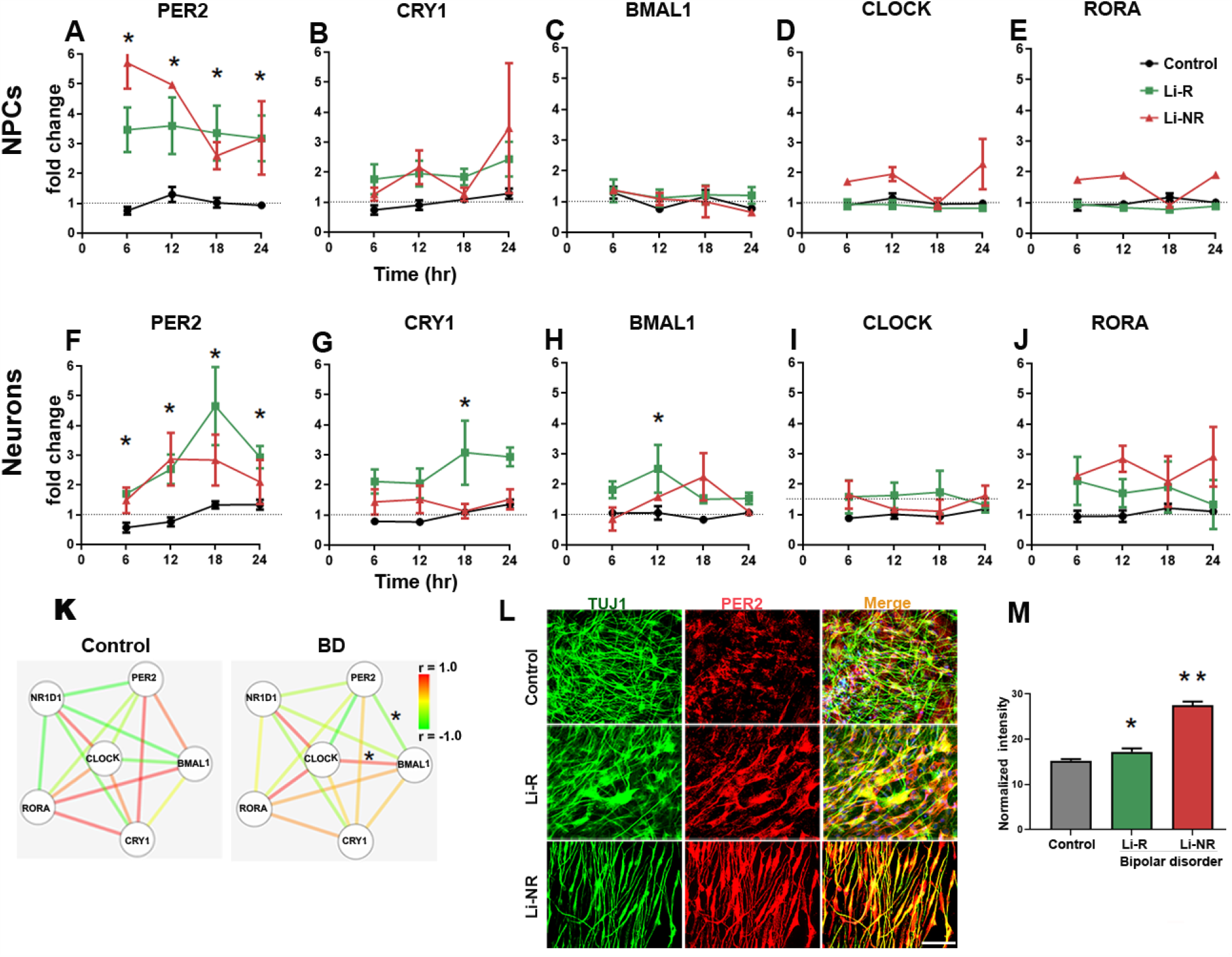
Circadian clock gene expression in NPCs and neurons. Results of RT-PCR 24 h time course experiments performed on mRNA collected at 6 h intervals in NPCs (A-E) and neurons (F-J). Data from controls (black), Li-R (green) and Li-NR (red) are shown. Results were analyzed with two-way ANOVA: * indicates a main effect of diagnosis in post-hoc test (p < 0.05). For each probe, expression was first normalized to *GAPDH*, and then to mean target expression over 24 h (dotted line). NPC means reflect controls: n=3 donors with 3-5 replicates/sample, Li-R: n=2 donors with 7-10 replicates/sample, Li-NR: n=2 donors with 7-10 replicates/sample. Neuron means reflect controls: n=3 donors with 4-5 replicates/sample, Li-R: n=2 donors with 7-10 replicates/sample, Li-NR: n=3 donors with 7-10 replicates/sample. All experiments were run in triplicate. (I) Clock gene network in BD and control neurons. Nodes indicate gene, edges indicate pairwise correlation. * indicates significant (p<0.05) difference in correlation coefficient (determined by two-way ANOVA with post-hoc test). (L-M) PER2 protein expression in control, Li-R and Li-NR neurons. PER2 expression was significantly higher in both groups of BD neurons (mean immunoreactivity in arbitrary units) ± SEM: control: 15.26 ± 0.34, Li-R 17.24 ± 0.72, Li-NR 27.47± 0.88, One-way ANOVA revealed F=112, p<0.0001, data reflect results from n= 350 control, 220 Li-R, 160 Li-NR neurons (quantified by DAPI staining), from n=3 control, 2 Li-R and 3 Li-NR donors). Post-test revealed * p<0.05 vs control, **p<0.001 vs control and Li-R. Error bars indicate standard error of the mean (SEM).

### Effects of lithium on neuronal rhythms

We treated neurons with lithium to identify effects on circadian rhythms. Lithium lengthened period in a concentration- and diagnosis-dependent manner, with significant effects on control and Li-R cells at 10 mM, but with no effect on Li-NR cells regardless of concentration (Figure 5A, by 2-way ANOVA: diagnosis (p < 0.002), drug (p < 0.02). Consistent with previous observations in fibroblasts ^12^, lithium caused a modest increase in amplitude controls at 1mM and decrease at 10 mM, but no significant effect on amplitude in BD neurons from either Li-R or Li-NR. Overall, this pattern of amplitude change reflected a trend-level difference in groups and a significant lithium x diagnosis interaction (Figure 5B).

**Figure 5:**
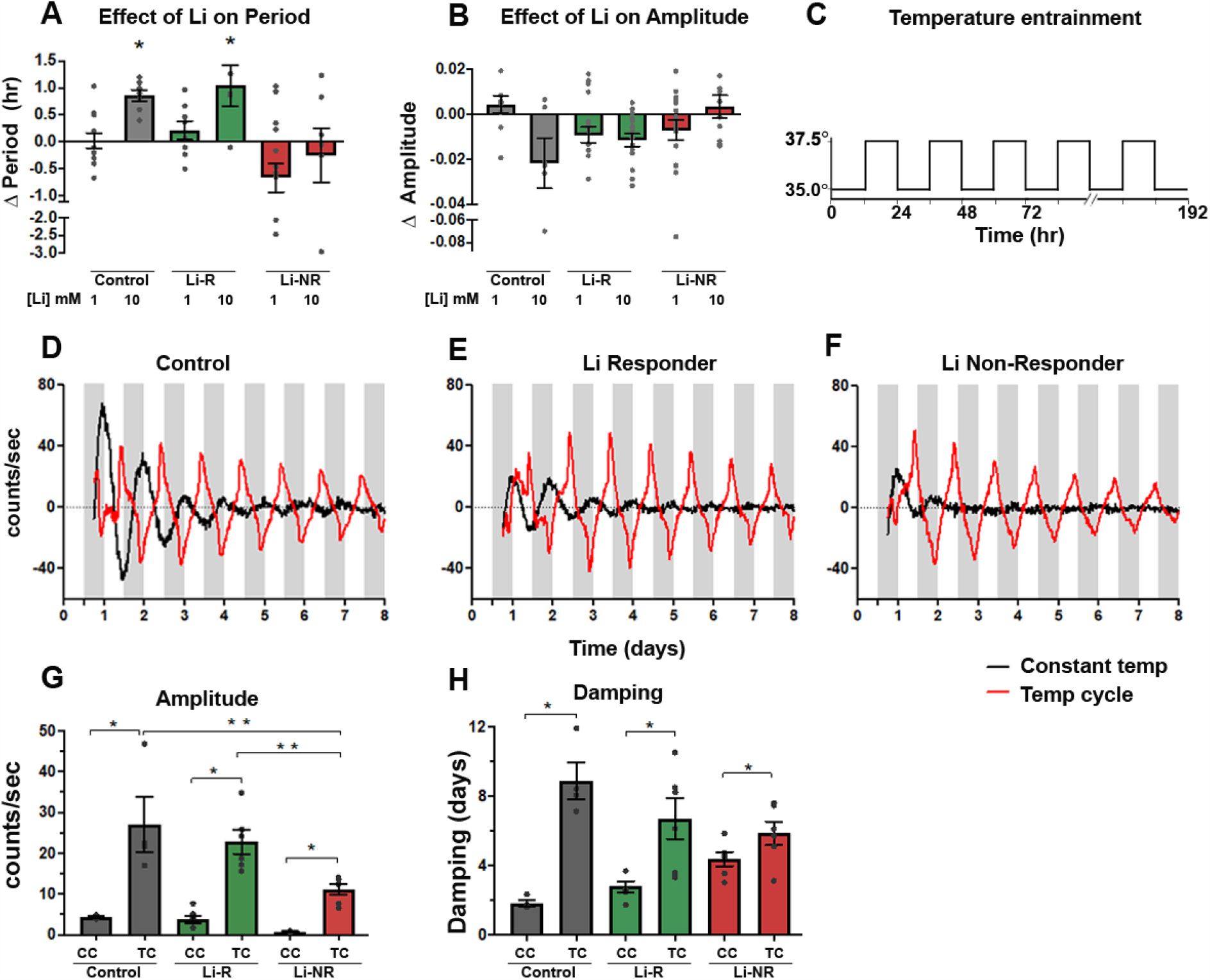
Effects of lithium and temperature entrainment on neuronal rhythms. Neurons were treated *in vitro* with lithium (1-10 mM) or vehicle **(A)** Effect of lithium on period. Lithium had a significant period-lengthening effect in controls and Li-R but not Li-NR (two-way ANOVA revealed effects of diagnosis (p<0.001) and diagnosis x lithium (p<0.02). **(B)** Effect of lithium on amplitude. Two-way ANOVA revealed a significant BD x lithium interaction effect on amplitude (p<0.01), whereby amplitude changes across lithium exposures differed significantly depending on diagnosis. Results **(A-B)** reflect the mean within-sample difference in period (lithium-vehicle) matched for experimental run for samples from n=3 controls, 2 Li-R, and 3 Li-NR averaged over three technical replicates in five separate experimental trials. **(C)** Schematic illustration of 12h/12h temperature entrainment schedule **(D-F)** Neurons from the same donor were studied in parallel under constant (CC) or temperature cycling (TC) conditions. Representative traces of *Per2-luc* rhythms are shown from control, Li-R, and Li-NR neurons, under CC (black) or TC (red). For the TC condition, white bars indicate 35°C, gray bars indicate 37.5°C. (G) Effects of TC on rhythm amplitude. Amplitude increased in the TC condition for all samples, but the increase was not as great in Li-NR neurons [2-way ANOVA: temperature (p<0.0001), diagnosis (p<0.02), diagnosis x temperature (p < 0.05)]. (H) Effect of TC on rhythm damping. Rhythms damped more slowly in the TC condition for all neurons, but this effect was not great in Li-NR neurons [2-way ANOVA: temperature (p<0.0001), diagnosis x temperature (p < 0.05)]. * indicates significant effect of TC, ** indicates significant group difference. Error bars indicate standard error of the mean (SEM).

### Effects of temperature entrainment on neuronal rhythms

Previous work has revealed rhythm differences in BD fibroblasts under temperature-entrainment ^13^. Therefore, we recorded rhythms in identically-prepared neurons (grown from n=1 control, 1 Li-R and 1 Li-NR) either at a constant 35°C (CC), or in parallel using a 12h:12h temperature cycle (TC, 35°C/37.5°C, Figure 5C). Across all groups, amplitudes under TC was significantly increased compared to CC. However, despite the increase, amplitude in the Li-NR samples remained lower than controls and Li-R (Figure 5G). Damping was slowed by TC, but the effect was only significant in control and Li-R neurons (Figure 5H).

## Discussion

Circadian rhythm disruption has been reported in BD for decades ^6, 21, 22^, and recent genetic studies more clearly demonstrate the overlap between the circadian clock and BD ^23-27^. Our work adds to this literature by directly demonstrating disrupted circadian rhythms in living NPCs and neurons from BD patients. In *Per2-luc* assays, we identified greatly reduced rhythm amplitude and faster damping in cells from Li-NR compared to control and Li-R cells. In NPCs, rhythm period was shorter in Li-R compared to Li-NR and controls. Single-cell analyses indicated reductions of rhythms and misalignment of phase in BD NPC and neurons. *Per2-luc* assays reflect the overall activity of the circadian network and are good for resolving differences in cellular rhythms, but cannot discern the varying contributions of underlying network components. To provide additional mechanistic insight, RT-PCR and protein assays were performed and showed increased PER2 expression, especially in Li-NR. As a transcriptional repressor, PER2 provides negative feedback to the circadian network and suppresses rhythms. Its proper function relies upon precise timing of protein turnover to regulate its inhibitory actions. Therefore, amplitude reductions and phase dispersal observed in rhythm assays may reflect the consequences of PER2 overexpression, leading to heightened negative feedback from PER2. The Li-R and Li-NR BD groups differed in important ways, with Li-R showing higher levels of neuronal *CRY1* and *BMAL1* gene expression, and Li-NR showing the highest levels of neuronal PER2, possibly underlying the more severe disruption of rhythms observed in Li-NR cells.

Consistent with previous findings ^12^, BD neurons show a differential sensitivity to the period-lengthening and amplitude-increasing effects of lithium. The response of neurons to the period-lengthening effect correlated with the donor’s clinical lithium response, suggesting overlap across therapeutic and circadian mechanisms ^8^. Interestingly, temperature entrainment partially reversed the low amplitude phenotype in Li-NR neurons suggesting that despite the loss of rhythm, some underlying core clock mechanisms remain intact in BD neurons and are amenable to intervention ^28^. These data imply that in BD, defects in the circadian clock may not lie entirely within the core-clock network, but perhaps also in regulatory inputs. This interpretation is consistent with fibroblast^29, 30^ and GWAS^27^ data in BD that implicate ion channels and entrainment-related signaling networks^31, 32^. Models of BD using iPSC-derived hippocampal neurons identified abnormalities in *KCNC1*/*KCNC2*-encoded potassium channels, causing unstable and hyperexcitable electrical activity, particularly in Li-NR^14, 33-35^. These findings are compatible with ours, as these channels have important roles in neuronal circadian rhythms^36^.

Our previous work on cellular rhythms in BD used fibroblasts. While the circadian clock is cell-autonomous and fully functional in fibroblasts^19^, they lack many essential neuronal features. Therefore, we developed an iPSC-based model of excitatory cortical neurons. Postmortem and MRI studies have indicated a loss of cortical volume in BD^37, 38^, with disturbances of the glutamate system ^39, 40^, supporting the study of these neurons in cellular models ^39^. Previous neuronal models of BD have used cortex-like cells to reveal lithium-sensitive differences in dendritic organization related to CRMP2 phosphorylation^41^. While the SCN is the master clock, clock gene expression in widespread throughout the primate brain^42^ and loss of cortical rhythms has been demonstrated in other psychiatric disorders such as major depression ^43^ and schizophrenia ^44^.

Previous data from BD patient fibroblasts have shown differences in period length and less sensitivity to the effects of lithium on rhythms ^12^, lower amplitude rhythms under temperature-entrained conditions ^13^ and an association of shorter period with Li-R ^8^. Our neuronal models replicate some aspects of these data and reveal interesting differences. Previously identified rhythm disruptions in BD fibroblasts were modest, rarely demonstrating a total lack of rhythms ^8, 12^. Compared to fibroblasts, Li-NR rhythms were more profoundly affected in neuronal cells, indicating BD-associated circadian disruption is more pronounced in this context. The single-cell analyses also mark an important advance over previous work. While low amplitude has been reported previously in BD fibroblasts^13^, it was unclear if the underlying mechanism was related to weak expression of the clock, phase desynchrony or some combination, as studies of cell populations cannot resolve these differences. In the single-cell studies of BD, we found evidence of both overall weaker rhythms and phase dispersal in NPCs and neurons.

In line with previous findings in peripheral cells ^8^, we found shorter circadian periods in NPCs from Li-R compared to Li-NR. That this effect was limited to NPCs and not observed in neurons may reflect a limitation of statistical power or could reveal a developmental interaction in NPCs between the clock and lithium’s therapeutic action. NPCs represent an immature, and mitotically active population that supports neurogenesis ^45^. Adult neurogenesis in the dentate gyrus may play a role in mood regulation, is promoted by lithium ^46, 47^ and has been linked to *Per2* and *Bmal1* ^48^. It is therefore interesting that the shorter period in Li-R relates specifically to mitotically active NPCs, suggesting the possibility of a circadian clock-neurogenesis link.

Another insight from our neuronal model relates to the role of development. In single cells, we observed that control neurons were more likely to be rhythmic than NPCs. This difference was absent in BD neurons, possibly suggesting developmental immaturity of the BD clock. Previous neuronal models of BD have implicated neurodevelopmental pathways ^49^, but the interaction between development and the circadian clock remains poorly described. Others have reported that and that the clock is developmentally regulated and active in embryonic stem cells, and that *Bmal1* plays a role in stem cell differentiation in mice ^50, 51^. However, no studies have investigated clock development in human neuronal models of neuropsychiatric disorders.

The strengths of our study are the novel use of a neuronal model to study circadian rhythms, and convergent methods to characterize rhythms, allowing us to report in detail the cellular, molecular, and temporal features of the BD circadian phenotype. Our BD subjects were all characterized for lithium response by prospective, long-term assessment during a monotherapy trial allowing clinical phenotypes to be determined with particularly high confidence ^5^. Our study also has limitations. While we used a protocol shown previously to generate electrically active neurons from iPSCs ^16^, we did not characterize electrical activity in this study and relied upon protein markers to identify neurons. Furthermore, we only studied glutamatergic neurons, and excluded other cell lineages worthy of study including serotonergic ^52, 53^, dopaminergic ^54^ and GABAergic ^55^ neurons. The study sample was small and relied upon sampling BD cases representing the extremes of clinical response. Since lithium response exists on a spectrum, our results may not apply in all cases. Moreover, we only had sufficient statistical power to identify large effects, and our methodology relied upon repeated sampling of a small number of subjects and therefore may over-estimate group differences. NPC data reflect the results of up to four controls, two Li-R, and three Li-NR cell lines. Neuron data reflect the means of up to three controls, two Li-R, and three Li-NR cell lines. To overcome these limitations, we made use of within-subject comparisons to reduce variability in experiments (e.g. comparing the same cell to itself after various treatments) and single-cell studies that allow for large numbers of cells to be studied. We are re-assured that many of our results are consistent with studies from fibroblasts using larger, independent cohorts ^8, 12, 13^. Nonetheless, replication of these results in larger cohorts will be essential.

In summary, we have shown for the first time that there are substantially disrupted circadian rhythms in live neurons from BD patients. The disruptions are most severe in Li-NR cells, but are also observed in Li-R in single-cell studies. Overexpression of *PER2* and negative transcriptional regulators that are core components of the molecular circadian clock may be an important contributor to this disruption, but some contributing factors likely occur outside the core-clock network and may be amenable to correction by targeted manipulation of inputs. Consideration of circadian timing opens a new dimension of analysis in neuronal models of BD and related psychiatric disorders which have previously been limited to cross-sectional analyses at a single time point ^14, 33^. In any given cell type, ∼10% of genes are rhythmically expressed ^56^. Given the rhythm disruption in BD neurons, a substantial number of clock-output genes are likely affected and may impact upon neuronal excitability and development, dimensions of BD pathophysiology that were previously identified in neurons. Rhythm abnormalities may serve as a cell-based biomarker and drug development platform to identify new molecular targets and pharmacological modulators of cellular rhythms in BD patient neurons. Future work in these areas should consider circadian timing.

## Acknowledgements

The authors wish to thank Anna Nilsson, Lucia Illescas, Rita Fischer, for technical assistance; and Carol Marchetto and Krishna Vadodaria for helpful input. The work was funded by VA Merit Awards to MJM (BX003431) and DKW (BX001146) and a Catalyst Research Award from the International Bipolar Foundation to MJM. Sample collection and clinical assessment in the PGBD Trial was supported by NIMH/NIGMS (MH92758), The Western Norway Regional Health Authority and the Canadian Institutes of Health Research (#64410).

