## Supplementary material for "Circadian rhythms in bipolar disorder patient-derived neurons predict lithium response"

**Supplementary Methods:**

**Pharmacogenetics of Bipolar disorder trial (PGBD**): Clinical response to lithium was determined prospectively in patients with BD type I between January 2011 and January 2016 through the PGBD multi-center treatment trial. After providing informed consent, phenotypic information on chronotype and psychiatric history was obtained at baseline. Over a 12-week period, subjects were then transitioned from their current medication regimen to lithium monotherapy. During this time, subjects were evaluated by a psychiatrist every 2 weeks and completed mood ratings. BD patients who were stabilized on lithium monotherapy were termed lithium responders (Li-R) while those who were unable to stabilize were classified as lithium non-responders (Li-NR). Subjects who left the study for intolerable side effects or reasons unrelated to their clinical response to lithium were excluded from further analysis. After 12 weeks, Li-R subjects entered a maintenance phase and were evaluated by the study physician in person every two months for up to 2-yr. Subjects were determined to have relapsed during maintenance if they reported a DSM-IVTR defined manic or depressive episode.

**Supplementary Tables:**

| **Cell ID** | **Age** | **Sex** | **Race** | **Diagnosis** | **Li Response** | **Days Stable Li** | **Morningness (BALM)** |
|---|---|---|---|---|---|---|---|
| C1 | 56 | M | C | Control | | | |
| C2 | 73 | M | C | Control | | | |
| C3 | 38 | M | C | Control | | | |
| C4 | 33 | M | C | Control | | | |
| BD1 | 57 | M | C | Bipolar | R | >730* | 48 |
| BD2 | 65 | M | C | Bipolar | R | >730* | 45 |
| BD3 | 69 | M | C | Bipolar | NR | 138 | 51 |
| BD4 | 22 | M | C | Bipolar | NR | 28 | 34 |
| BD5 | 54 | M | C | Bipolar | NR | 88 | 28 |

**Table S1. Clinical Characteristics of cell line donors.** * indicates stable throughout entire duration of 2-year study. BALM: Basic language morningness scale. Identification numbers were assigned arbitrarily and applied consistently to the same donor across all tables and figures.

| Target | Species | Company | Cat # | Dilution |
|---|---|---|---|---|
| TUJ-1 | Chicken | Biolegend | 801201 | 1:300 |
| GFAP | Goat | Abcam | ab53554 | 1:600 |
| NESTIN | Mouse | Millipore | MAB5326 | 1:200 |
| SOX2 | Rabbit | Cell Signalling Tech | 3579S | 1:300 |
| TRA 1-60 | Mouse | Millipore | MAB4360 | 1:300 |
| NANOG | Goat | R&D Systems | AF1997-SP | 1:400 |
| VGLUT2 | mouse | Synaptic Systems | #135 421 | 1:300 |
| PER2 | Rabbit | Abcam | ab179813 | 1:300 |

**Table S2. List of primary antibodies**

| | Marker ICC | Lumino-meter | Single cell | qPCR | Lithium |
|---|---|---|---|---|---|
| C1 | Y | Y | | Y | Y |
| C2 | Y | Y | Y | Y | Y |
| C3 | Y | Y | Y | Y | Y |
| C4 | | Y | | | |
| BD1 (Li-R) | Y | Y | Y | Y | Y |
| BD2 (Li-R) | Y | Y | Y | Y | Y |
| BD3 (Li-NR) | Y | Y | | Y | Y |
| BD4 (Li-NR) | Y | Y | | Y | Y |
| BD5 (Li-NR) | Y | Y | Y | | Y |

**Table S3. NPC lines used in experiments.** Marker: NESTIN/SOX2, qPCR: Time course using real time quantitative polymerase chain reaction, ICC: immunocytochemistry. C: control, BD: bipolar, Li-R: lithium responder, BD Li-NR: lithium non responder. Identification numbers correspond to phenotypic information reported in Table S1.

| | Marker ICC | Lumino-meter | Single-cell | qPCR | PER2 ICC | Lithium | T-Cycle |
|---|---|---|---|---|---|---|---|
| C1 | Y | Y | Y | Y | Y | Y | Y |
| C2 | Y | Y | Y | Y | Y | Y | |
| C3 | Y | Y | Y | Y | Y | Y | |
| C4 | | | | | | | |
| BD1 (Li-R) | Y | Y | Y | Y | Y | Y | Y |
| BD2 (Li-R) | Y | Y | Y | Y | Y | Y | |
| BD3 (Li-NR) | Y | Y | Y | Y | Y | Y | Y |
| BD4 (Li-NR) | Y | Y | | Y | Y | Y | |
| BD5 (Li-NR) | Y | Y | | Y | Y | Y | |

**Table S4. Neuron lines used in experiments.** Marker: TUJ1/GFAP/VGLUT2, qPCR: Time course using real time quantitative polymerase chain reaction, ICC: immunocytochemistry. C: control, BD: bipolar, Li-R: lithium responder, BD Li-NR: lithium non-responder. Identification numbers correspond to phenotypic information reported in Table S1.

### Supplementary Figures

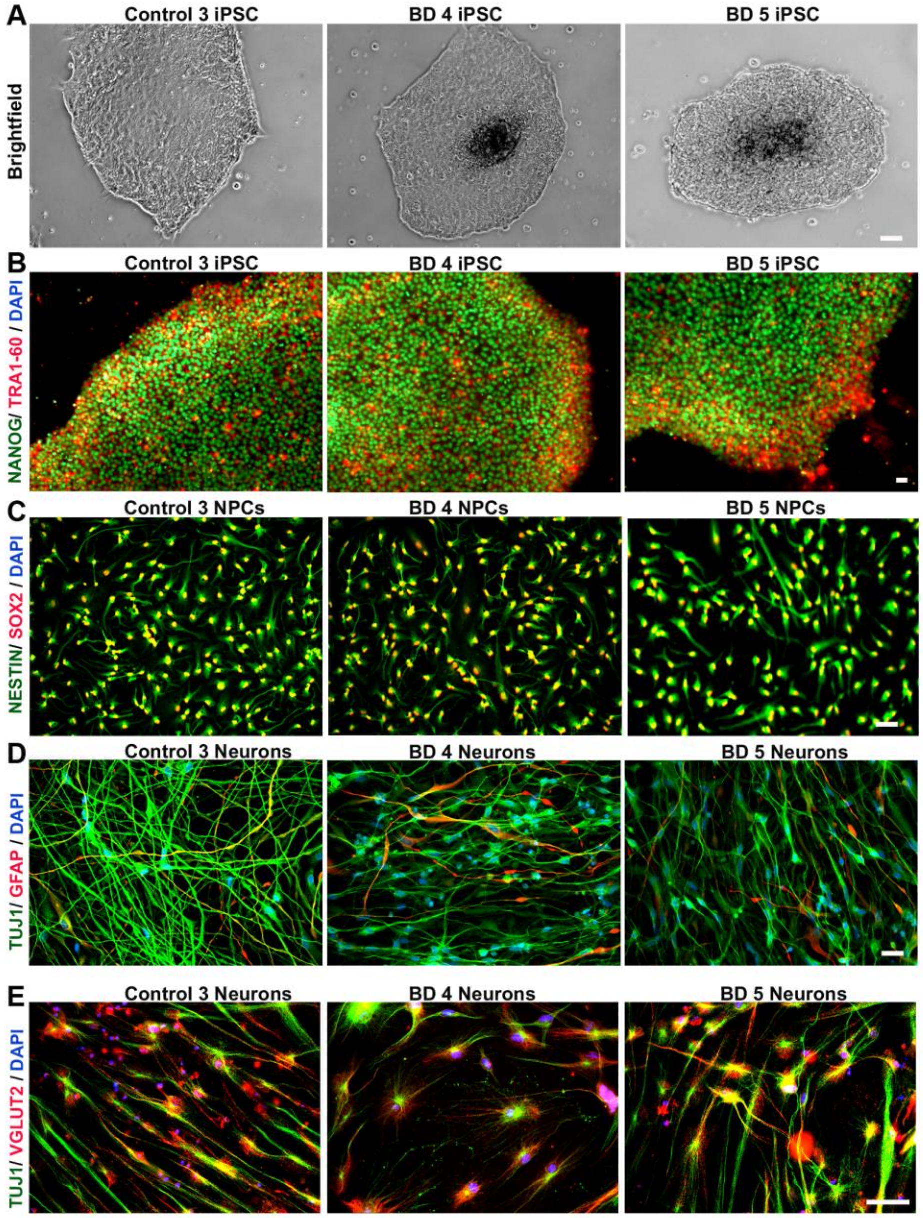


**Figure S1: Generation of human neurons from induced pluripotent stem cells**. Additional examples of (A) Brightfield images of iPSC clones demonstrating morphological features (B) Representative images of iPSC clones from control and BD patients co-labeled with the pluripotency markers, TRA 1-60 (red), and NANOG (green). Cell nuclei are stained with DAPI. (C) Representative images of NPCs from control and BD donors showing co-expression of early cortical neural precursor markers NESTIN (green) and SOX-2 (red). (D) Differentiated neurons expressing TUJ1 (neuron-specific) and GFAP (glia-specific) markers from control and BD cell lines. (E) Representative images of control and BD neurons showing cellular co-localization of vesicular-glutamate transporter 2 (VGLUT2) and TUJ1. Nuclei were visualized with DAPI. Scale bars = 50µm. BD: bipolar disorder. Identification numbers correspond to subject data reported in Tables S1, S3, S4.

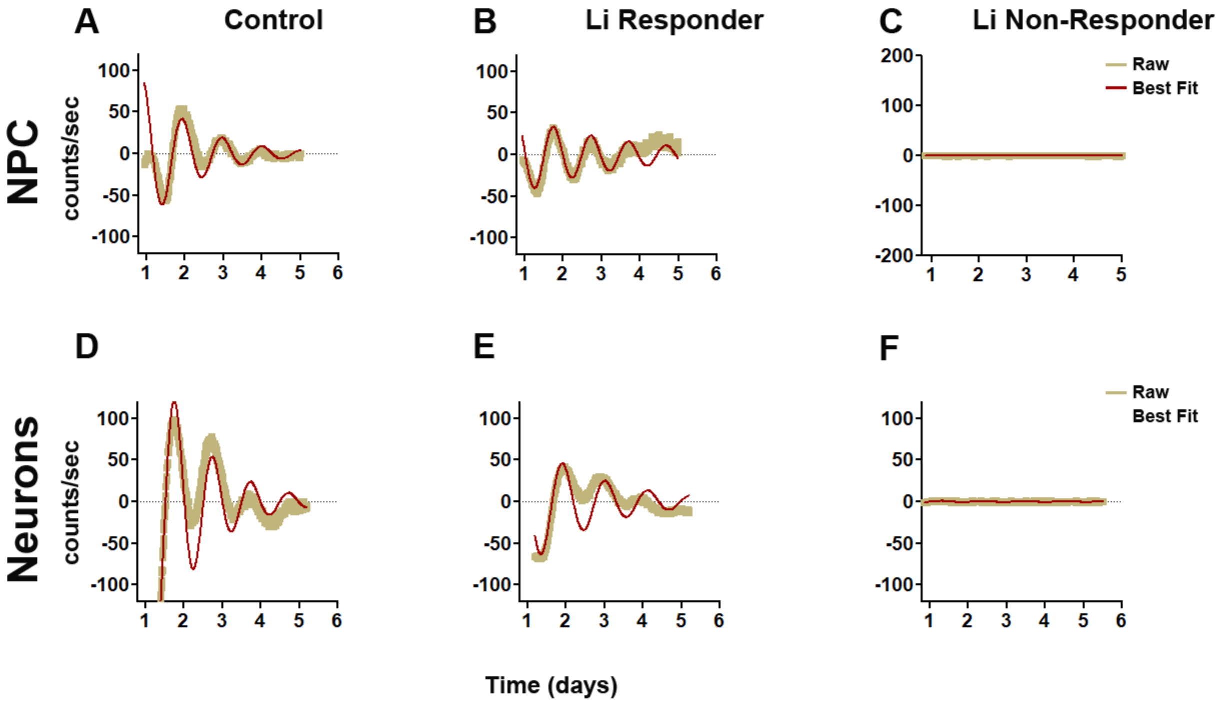


**Figure S2: Luminometer assays of circadian rhythms in NPC and neurons.** Representative rhythm traces from additional cell lines showing Per2-luc expression measured in NPCs from **(A)** control (**B**) lithium responders (Li-R), and **(C)** lithium non-responders (Li-NR). Yellow indicates raw counts, red indicates best fit line. Representative traces of Per2-luc rhythms measured in neurons from **(D)** control (**E**) lithium responders (Li-R), and **(F)** lithium non-responders (Li-NR). NPC data reflect the results of n=4 controls, 2 Li-R, and 3 Li-NR cell lines, recorded in triplicate. Neurons data reflect the results of n=3 controls, 2 Li-R, and 3 Li-NR cell lines, recorded in triplicate.

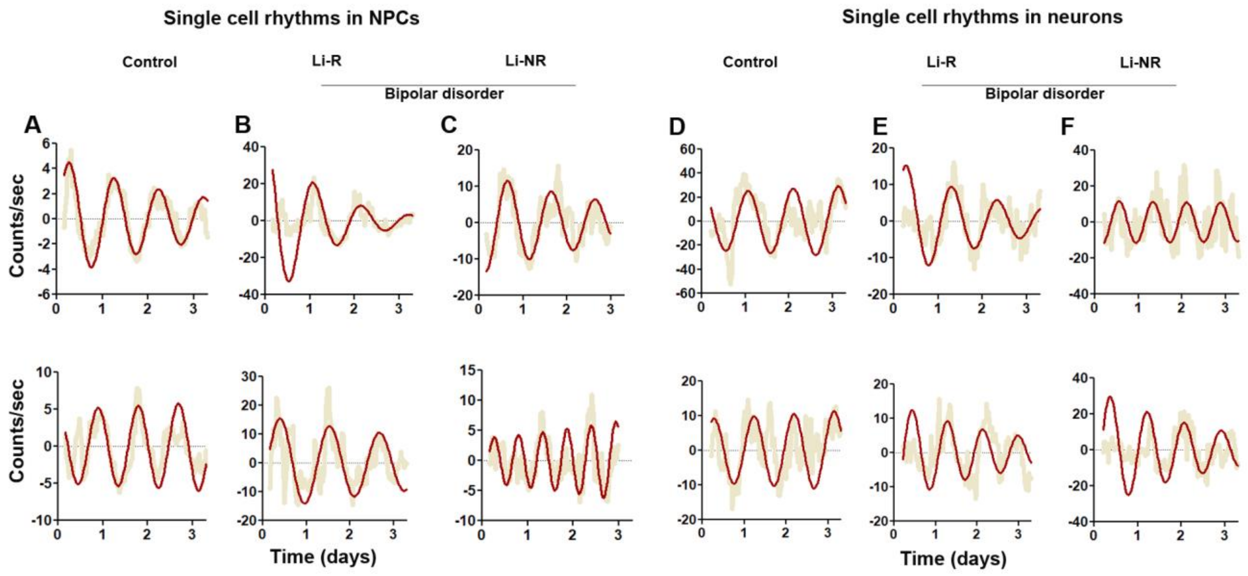


**Figure S3: Single cell assays of circadian rhythms in NPC and neurons.** Representative rhythm traces from additional single cells showing Per2-luc single cell expression in NPCs from **(A)** control **(B)** lithium responders (Li-R), and **(C)** lithium non-responders (Li-NR). Yellow indicates raw counts, red indicates best fit line. Representative traces of Per2-luc single cell rhythms measured in neurons from **(D)** control **(E)** lithium responders (Li-R), and **(F)** lithium non-responders (Li-NR). Unlike the data shown in Figure 3, cells are depicted irrespective of phase. NPC data reflect the results of single cell experiments of 60-80 cells from n=2 controls, 2 Li-R, and 1 Li-NR donors. Neuron data reflect the results of 60-80 cells from n=3 controls, 2 Li-R, and 1 Li-NR donors.

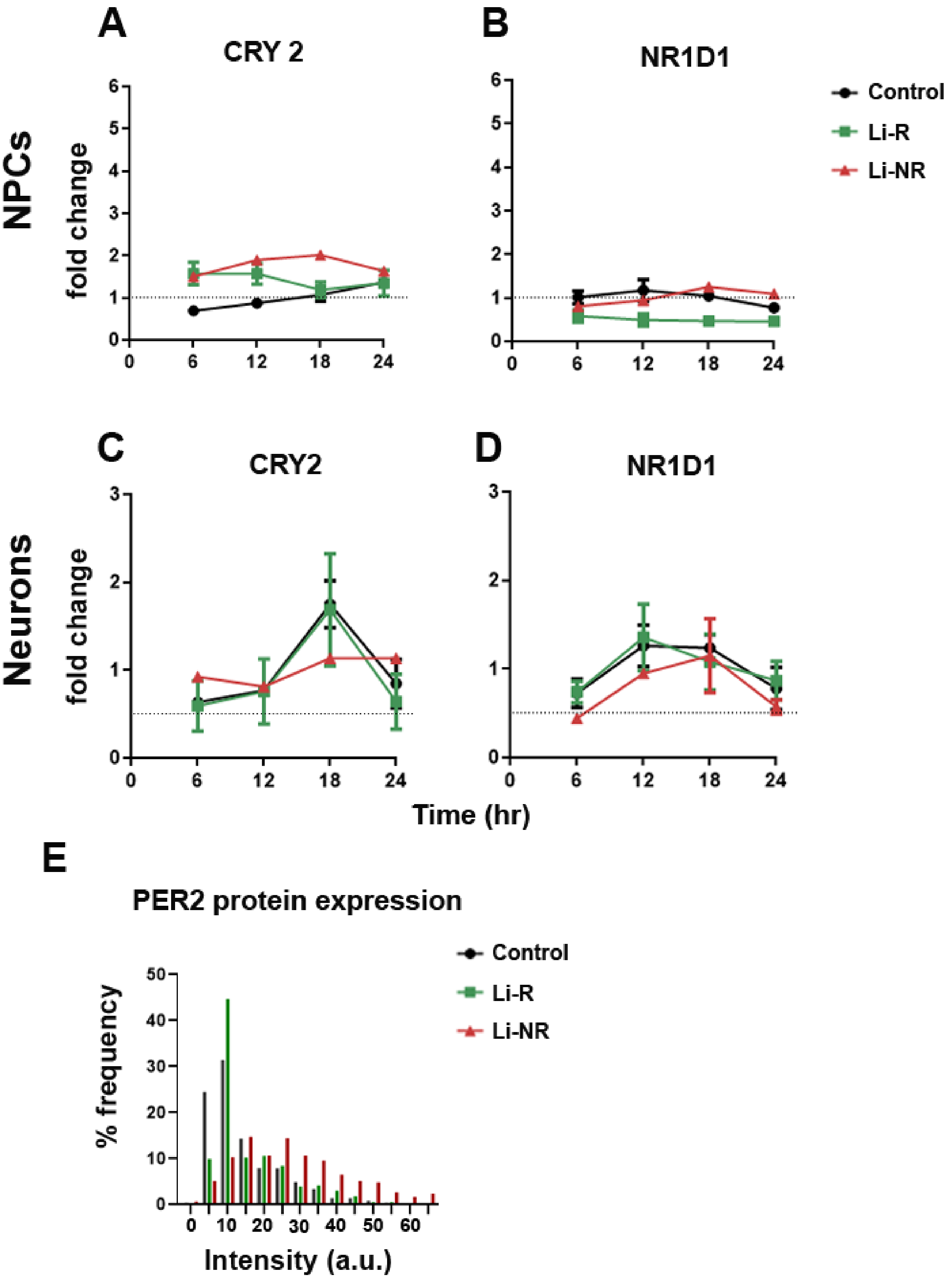
A
CRY 2
B
NR1D1
Control
Li-R
Li-NR
NPCs
fold change
C
CRY2
D
NR1D1
Neurons
fold change
Time (hr)
E
PER2 protein expression
Control
Li-R
Li-NR
% frequency
Intensity (a.u.)

**Figure S4: Circadian clock gene expression in NPCs and neurons.** Additional data for *CRY2* and *NR1D1* expression in NPCs (A, B) and neurons (C, D). Control data are shown in black and BD is shown in red. NPC results reflect controls: n=2 donors with 3-5 replicates/sample, BD: n=3 donors with 7-10 replicates/sample. Neuron results reflect controls: n=2 donors with 4-5 replicates/sample, BD: n=5 donors with 7-10 replicates/sample. All replicates were run in triplicate. None of the differences in A-D were statistically significant. Error bars indicate standard error of the mean (SEM). (E) Frequency distribution of PER2 intensity in the total population of control, Li-R and Li-NR neurons (n=3 control, 2 Li-R, and 3 Li-NR). Intensity was normally distributed. Low intensity staining was enriched in control and higher intensity staining was significantly enriched in BD samples, especially in the Li-NR neurons.
